# CD2AP’s Structure and Oligomerization are compromised by the K301M mutation: implications for Nephrotic syndrome

**DOI:** 10.64898/2026.01.08.698362

**Authors:** Abrar H Qadri, Raja Mani Tripathi, Iqball Faheem, Samiksha Meshram, Eswar R. Reddem, Ravishankar Ramachandran, Anil K Pasupulati

## Abstract

**Introduction:** The podocyte slit diaphragm (SD) is a complex filtration unit localized to the blood and urine interface and governs the glomerular selectivity. However, the greater details of the SD composition and the mechanism of assembly of the SD protein as a macromolecular complex remain elusive. CD2-associated protein (CD2AP) serves as a central scaffold within the SD, and mutations in CD2AP are strongly associated with nephrotic syndrome (NS) and focal segmental glomerulosclerosis (FSGS). However, the mechanisms by which such mutations alter the architecture and higher-order organization of CD2AP are poorly understood.

**Methods:** We employed biophysical, structural, and proteomic approaches to investigate the impact of the disease-associated K301M mutation on CD2AP structure and its interaction with Podocin. Oligomerization was analyzed using size-exclusion chromatography, blue native PAGE, Dynamic light scattering, and small-angle X-ray scattering. Secondary and tertiary structural properties were assessed by far- and near-UV circular dichroism, thermal denaturation, and intrinsic fluorescence spectroscopy. CD2AP–podocin interactions were quantified using in vitro pulldown and surface plasmon resonance (SPR), and mutation-dependent changes in interaction networks were examined through interactome profiling.

**Results:** Wild-type (WT) CD2AP assembled into flexible higher-order oligomers (∼9–12-mers), whereas the K301M variant collapsed into lower-order species (∼3–6-mers), indicating destabilization of the coiled-coil assembly interface. Spectroscopic analyses revealed subtle secondary-structure rearrangements, but profound tertiary packing defects, as well as reduced and markedly diminished thermal resilience in the mutant. SPR analysis demonstrated loss of binding between Podocin and mutant CD2AP, whereas WT CD2AP showed high-affinity interaction (KD = 211 nM) with Podocin. Complementary interactome profiling revealed widespread rewiring of protein–protein interactions in the case of mutant CD2AP, characterized by the loss of core partners and the emergence of aberrant associations.

**Conclusion:** These findings define a mechanistic model in which the K301M mutation destabilizes CD2AP oligomerization, disrupts podocin recognition, and remodels interaction networks essential for SD stability. This work signifies the importance of CD2AP in SD assembly and the permselective filtration function of the kidney, and the impact of a single mutation in the pathogenesis of NS and FSGS.

**Translational Statement:** Inherited nephrotic syndrome, characterized by heavy proteinuria, frequently arises from mutations in scaffolding proteins of the slit-diaphragm (SD). This study demonstrates that the nephrotic syndrome-associated K301M mutation in CD2-associated protein (CD2AP) compromises higher-order oligomerization, abolishes its binding to the binding partner (Podocin), and reshapes protein-protein interaction networks that are critical for SD assembly and stability. By establishing a direct link between mutation-induced collapse of CD2AP architecture and loss of podocyte scaffolding function, these findings provide mechanistic insight into the pathogenesis of CD2AP-associated proteinuric kidney disease. The results further identify oligomeric assembly interfaces as potential targets for therapeutic strategies aimed at preserving the integrity of the SD and glomerular filtration function.

Graphical Abstract

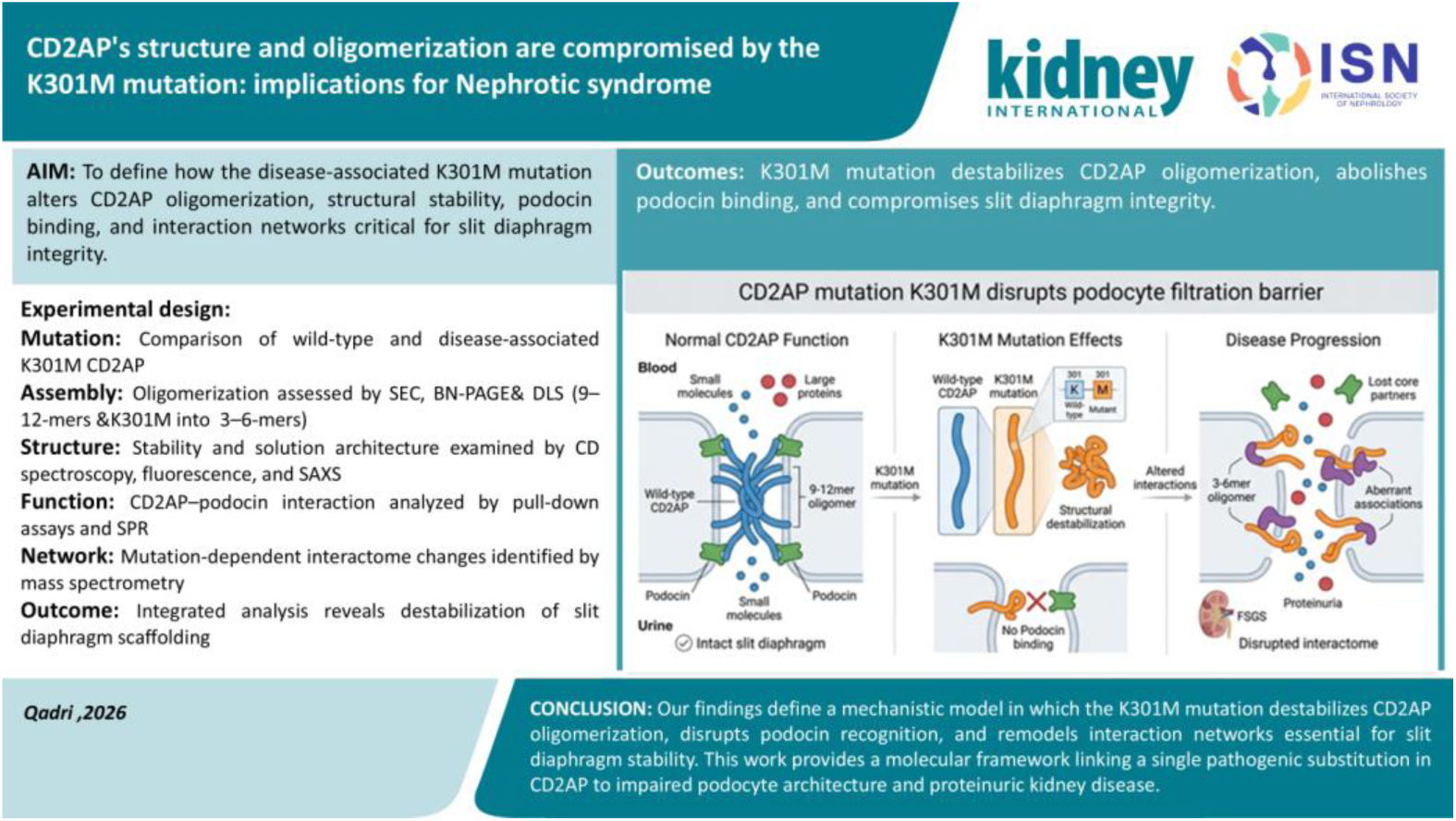

## Introduction

Nephrotic syndrome (NS) represents a primary clinical manifestation of glomerular injury characterized by massive proteinuria, hypoalbuminemia, and edema. Persistent proteinuria reflects dysfunction of the glomerular filtration barrier (GFB) and is an independent risk factor for the progression of chronic kidney disease (CKD) ^1, 2^. Corticosteroid therapy is effective for managing steroid-sensitive nephrotic syndrome (SSNS), but steroid-resistant nephrotic syndrome (SRNS) often advances to end-stage kidney disease (ESKD)^3^. A substantial portion of congenital and familial cases of NS have been linked to monogenic mutations in several podocyte-specific genes, which disrupt the permselectivity of GFB and cause proteinuric kidney diseases like Focal Segmental Glomerulosclerosis. Several of these mutations are associated with the proteins that constitute the podocyte (SD), which serves as a size- and charge-selective barrier to retain plasma proteins and filter out water and small molecules, forming part of the primary urine ^4^. SD, besides being a specialized cell-cell junction, also serves as a structural support and a signaling hub, thereby ensuring the integrity of the GFB^5^. The SD complex comprises nephrin, Podocin, CD2-Associated Protein (CD2AP), NEPH1, and transient receptor potential cation channel 6 (TRPC6) ^6–9^ ^10–14^. These proteins form an interconnected scaffold that connects the filtration slit to the actin cytoskeleton ^15–17^. This connection ensures mechanical stability and also integrates SD with intracellular signaling, which is essential for podocyte survival ^18, 19^. A preliminary structural insight provided by a recent cryo-electron tomography, where the SD components form a fishnet-like lattice generated by crisscrossing nephrin–NEPH1 heterodimers. However, the precise architecture and interaction network of SD at the molecular level remain largely elusive ^8, 20^.

Among SD proteins, CD2AP functions as a multidomain adaptor, anchoring the nephrin podocin complex to the actin cytoskeleton and thereby maintaining barrier resilience ^6, 10, 21, 22^. CD2AP comprises three Src homology 3 (SH3) domains, intrinsically disordered regions (IDRs), and a C-terminal coiled-coil domain that mediates oligomerization and multivalent interactions ^6, 10, 21–23^. Through these features, CD2AP coordinates cytoskeletal dynamics, endocytic trafficking, and signaling pathways that are essential for podocyte survival^15^.CD2AP-deficient mice exhibit congenital NS, characterized by podocyte effacement and early kidney failure ^3, 24–28^. The K301M substitution in CD2AP is associated with kidney disease; however, the molecular mechanism linking this mutation to NS remains poorly understood.

The molecular organization of the SD remains incompletely understood due to the intrinsic disorder and conformational flexibility of its components. Biophysical analyses of individual SD proteins could provide mechanistic insight into their assembly and cooperative interactions. Here, we present an integrated biophysical and structural characterization of wild-type CD2AP and its disease-causing variant, K301 M CD2AP. Our findings demonstrate that the K301M mutation disrupts CD2AP self-assembly and weakens its interaction with Podocin, thereby destabilizing the SD scaffold. These insights provide a molecular framework for understanding how pathogenic CD2AP variants impair podocyte integrity and for developing targeted strategies to preserve SD function and combat proteinuria.

## Materials and methods

### CD2AP expression and purification

A codon-optimized synthetic gene encoding full-length human CD2AP was obtained from Genscript and cloned into the pET-28a(+) vector to generate an N-terminal His₆-tagged construct. The K301M point mutation was introduced using overlap-extension PCR with gene-specific cloning primers (CD2AP-F1 and CD2AP-R1) and mutation-specific primers (CD2AP-K301M-F and CD2AP-K301M-R). The mutant construct was cloned into the same expression vector, and the presence of the K301M substitution was confirmed by Sanger sequencing. Plasmids encoding WT and K301M CD2AP were transformed into *E coli* Rosetta (DE3) competent cells for protein expression. Primer sequences are provided in **Supplementary Figure 1D.** Protein expression was induced with 1 mM IPTG at 16°C for 16 hrs. Cells were harvested by centrifugation at 8000 RPM for 20 min at 4°C and resuspended in lysis buffer A (10 mM potassium phosphate, 500 mM NaCl, 5 mM KCl 8% glycerol, pH 7.6), supplemented with 0.1% Triton X-100, a protease inhibitor cocktail, and 0.1 mg/ml lysozyme. Furthermore, after Sonication, cell debris was removed by centrifugation at 16,639 × g for 90 minutes at 4°C. The clarified lysate was loaded onto a 5 mL HisTrap HP column (Cytiva, Cat. No. 17524801) that was pre-equilibrated with buffer A containing 45 mM imidazole. The column was washed with buffer A supplemented with 100 mM imidazole, and the bound CD2AP protein was eluted using buffer A containing 400 mM imidazole. Eluted fractions were pooled and dialyzed against buffer A to remove imidazole. Protein concentrations were determined using a theoretical extinction coefficient of 43890 M⁻¹ cm⁻¹ (A280) and confirmed by the bicinchonic acid assay.

### Podocin Expression and purification

The codon-optimized human podocin gene (Genscript, Piscataway, NJ, USA) was cloned into the pMAL-p2x vector between the EcoRI and XhoI restriction sites to generate an N-terminal MBP fusion construct. Recombinant MBP–Podocin was expressed in *E. coli* Arctic Express (DE3) cells grown in LB medium at 37 °C and induced with IPTG. Cells were harvested by centrifugation and resuspended in lysis buffer (Buffer A: 20 mM sodium phosphate, pH 7.3, 0.5 M NaCl, 4.7 mM KCl, and 10% glycerol) supplemented with protease inhibitors, lysozyme, DNase I, and Triton X-100. After incubation on ice for 30 min, cells were disrupted by probe sonication, and insoluble material was removed by centrifugation at 10,000 × g for 1 h at 4 °C. The clarified lysate was applied to a pre-equilibrated amylose affinity prepacked column, a 5 mL MBP trap (Cytiva 28-9187), and washed with Buffer A containing 3 mM maltose. The column was then eluted stepwise with Buffer A supplemented with 35 mM maltose. Protein-containing fractions were analysed by SDS–PAGE, pooled, dialyzed against Buffer A, and quantified by absorbance at 280 nm (Jasco V-630) and BCA assay.

### Size Exclusion Chromatography

The oligomeric Nature of the WT and K301M CD2AP was determined by injecting 250 µL of 13.6 µM protein into a Superose 6 Increase 10/300 column (Cytiva) connected to an NGC system (Bio-Rad). The column was pre-equilibrated with 50 mM Tris-HCl, 300 mM NaCl, 4 mM EGTA, 5 mM KCl, and 7% glycerol, pH 7.6. The peak fractions were collected and analyzed on SDS-PAGE.

### Blue-Native Page (BN-PAGE)

BN-PAGE was used to corroborate the SEC results. SEC-purified CD2AP (30–40 μg) was loaded onto a 8% & 6% non-denaturing polyacrylamide gel and resolved at 4 °C. The gel system consisted of 1.5 M Tris-HCl (pH 8.8) with Coomassie G-250 added to impart a negative charge to the proteins. The cathode buffer consisted of 118 mM Tris-HCl, 39 mM glycine, and 0.05% G-250, while the anode buffer was composed of Tris-glycine (pH 8.0). The protein samples were mixed with 5X loading dye containing 20% glycerol, 0.2 M Tris-HCl (pH 6.8), 0.05% bromophenol blue, 1% Triton X-100, and 0.25% G-250. Gels were run at 150 V at 4°C and stained with Coomassie Brilliant Blue R-250.

### Circular Dichroism (CD) Spectroscopy

CD spectra of recombinant WT and K301M CD2AP were recorded on a JASCO J-1500 CD spectrophotometer equipped with a thermoelectric (Peltier) cell holder. For far-UV spectra (200–260 nm), a protein concentration of 5.46 µM was used in a 0.2 cm pathlength quartz cuvette, with a bandwidth of 2.5 nm and a scan speed of 50 nm/min. Near-UV spectra (260–360 nm) were recorded at 35 µM protein using a 0.5 cm pathlength cuvette and a scan speed of 100 nm/min. The spectra were recorded in triplicate and normalized against the buffer for both near- and far-UV CD measurements. Next, we analysed the effect of thermal-induced unfolding and the stability of the secondary structural elements in WT and K301M CD2AP. The protein samples were subjected to a steady increase in temperature, and spectra were recorded in the far-UV region at 10°C intervals using the same parameters.

### Intrinsic Tryptophan Fluorescence Spectroscopy

The intrinsic fluorescence spectra of WT and K301M CD2AP were recorded using a Jasco FP-6300 spectrofluorometer, equipped with a xenon flash lamp as the excitation source. Samples were excited at 286 nm, and emission spectra were collected over the 300–450 nm range. Spectra were acquired at various temperatures (20°C intervals), and emission intensities at 341 nm of both WT &K301M CD2AP were plotted to assess thermal denaturation. All measurements were triplicated using a 5 nm bandwidth and a scan speed of 100 nm/min. A protein concentration of 5.46 µM was used, and all spectra were buffer-subtracted.

### Dynamic Light Scattering (DLS)

Hydrodynamic size and aggregation state of WT and K301M CD2AP were determined using a DLS instrument. Proteins were dialyzed into 20 mM potassium phosphate buffer (pH 7.6), 300 mM NaCl, and 5% glycerol. All buffer components were filtered using 0.1 µm filters (Millipore). Measurements were first taken for the buffer, followed by samples containing 5.46 µM protein. Urea-induced unfolding was performed by incubating the protein with 8 M urea for 1 hour at room temperature. Data were analyzed using GraphPad Prism, and each experiment was conducted in at least two independent replicates.

### Small-Angle X-ray Scattering (SAXS)

SAXS data from solutions of CD2AP and CD2AP K301M point mutant in 10 mM potassium phosphate, pH 7.6 were collected on the Anton Paar SAXSpace instrument using a Mythen2 R 1K detector at a sample-detector distance of 0.3 m and at a wavelength of λ = 0.514 nm (I(s) vs s, where s = 4πsinθ/λ, and 2θ is the scattering angle). The solute concentration was measured at 10 °C to be 7.00 mg/ml. Four successive 900-second frames were collected. The data were normalized to the intensity of the transmitted beam and radially averaged; the scattering of the solvent blank was subtracted. Both datasets were deposited in SASDB (Small Angle Scattering Data Bank) with ID-SASDXJ6 and SASDXK6, respectively. 3D ab initio models were generated from SAXS data using DAMMIF analysis (ATSAS package). Chimera v1.15 was used to fit the model in the SAXS-generated envelope. Further data collection, structural, and modeling parameters are listed below.

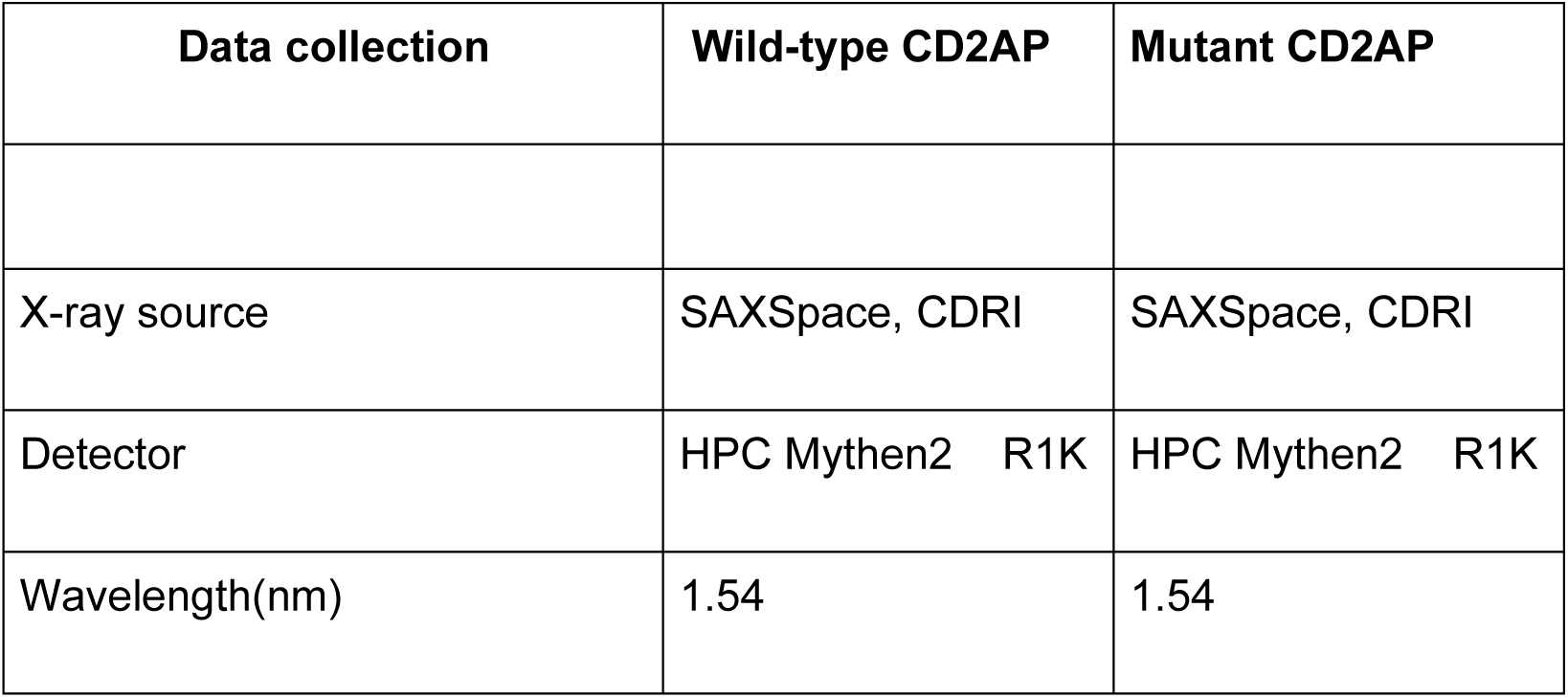
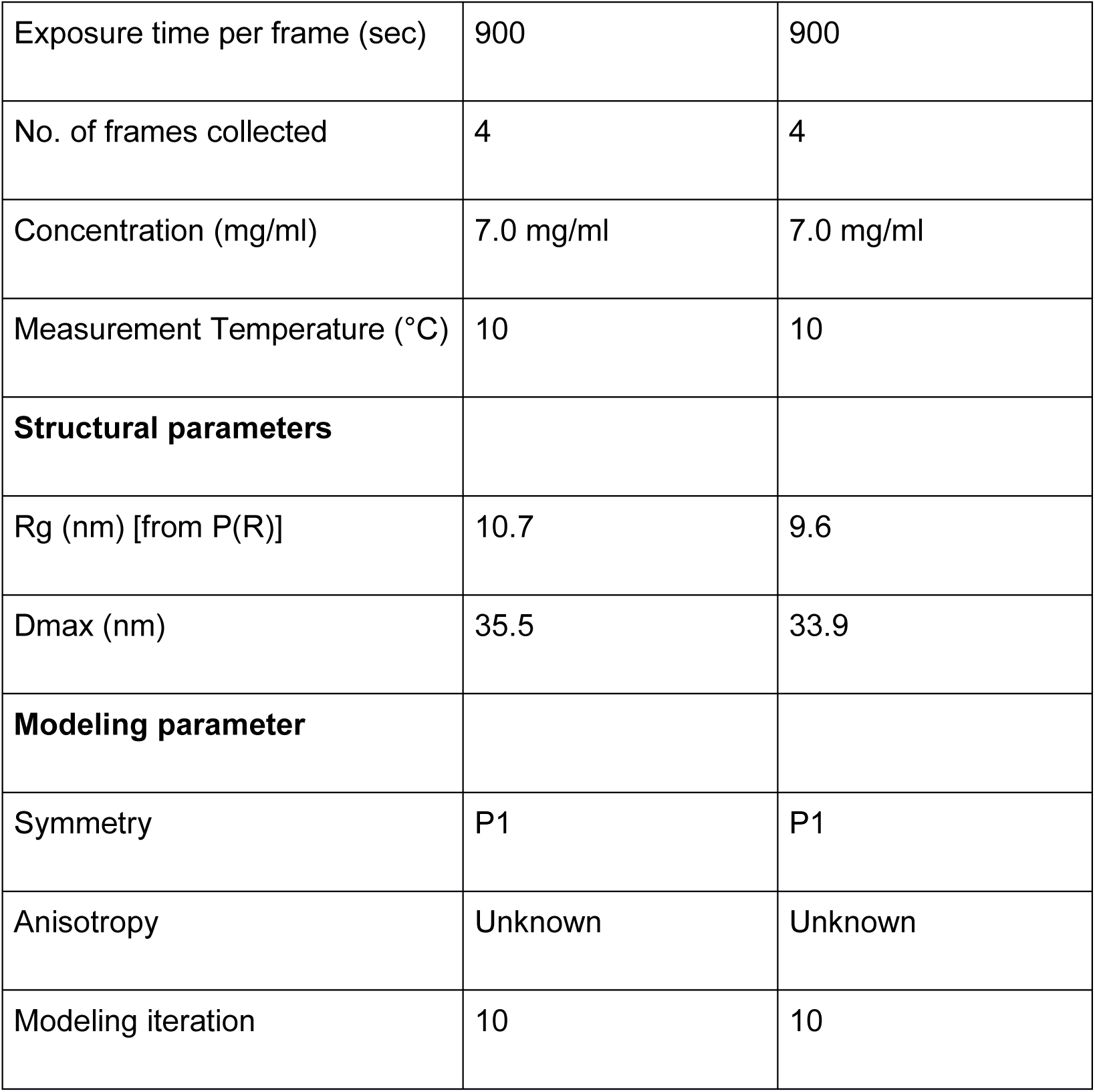

## In Vitro Pulldown Assays

### His-CD2AP as Bait

20–30 µg of His-tagged CD2AP was incubated with 50 µL of Ni-NTA resin (Qiagen) in binding buffer (20 mM Tris-HCl, 150 mM NaCl, 5% glycerol, 0.05% NP-40, pH 7.4) for 1 hour at 4 °C. After immobilization, 50 µg of MBP-Podocin was added and incubated for 2 more hours. The resin was washed five times with high-salt buffer (binding buffer with 300 mM NaCl), and SDS-PAGE and silver staining were used to visualize bound proteins.

### MBP-Podocin as Bait

Similarly, 50 µL of amylose resin was incubated with 20–30 µg of MBP-tagged Podocin, followed by the addition of 50 µg of His-CD2AP (WT or mutant). Further incubated for 2 hours, followed by washing five times with high salt buffer (binding buffer with 300 mM NaCl), and SDS-PAGE and silver staining were used to visualize bound proteins. Negative controls included resin-only and bait-only conditions.

### Surface Plasmon Resonance (SPR)

SPR assays were performed on a Biacore 3000 instrument (GE Healthcare) to quantify the interactions between CD2AP (WT and Mutant) and Podocin. Anti-His antibodies were immobilized on a CM5 chip via amine coupling (∼8000 RU). His-tagged WT or K301M CD2AP (3 µM) was captured on a CM5 chip (∼2000 RU from baseline), and Podocin was injected over the surface at serial dilutions (0 to 1.2 µM and 0 to 5µM) in HBS-EP buffer (10 mM HEPES, 150 mM NaCl, 3 mM EDTA, 0.005% Tween-20, pH 7.4) at 30 µL/min. The association between CD2AP and Podocin was monitored for 120 seconds. Binding kinetics and KD values were computed using BIA evaluation software by fitting to a 1:1 Langmuir binding model.

### In vitro pulldown and mass spectrometry-based interactome analysis

Purified WT and K301M mutant CD2AP proteins (200 µg each) were incubated with pre-washed and buffer-equilibrated Ni–NTA agarose beads at 4 °C for 2 hrs. to facilitate protein immobilization. Following incubation, the beads were thoroughly washed with binding buffer to remove unbound protein. Subsequently, human podocyte cell lysates were added to the beads and incubated at 4°C for 4 hrs (35 mM Tris–HCl (pH 7.4), 100 mM KCl, 0.5% NP-40, 5% glycerol) supplemented with protease inhibitor cocktail (Sigma Cat no 11873580001), to allow interaction with potential binding partners. Beads were washed with the same buffer as resuspension buffer, resuspended, and resolved on a 10% SDS–PAGE gel for visualization. The remaining bead-bound fractions were subjected to trypsin digestion for LC-MS/MS analysis using a protocol described in ^29^. The resulting peptides were purified using C18 reverse-phase columns and submitted for LC–MS/MS analysis to identify and compare the interactomes associated with WT and K301M CD2AP.

## Results

### Recombinant Purification of WT, K301M CD2AP and Podocin

Recombinant WT and K301M mutant human were heterologously expressed in *Escherichia coli* Rosetta (DE3) cells under conditions optimized to enhance soluble protein yield. Given the extensive intrinsically disordered regions present in CD2AP, expression parameters were systematically optimized. Induction with 1 mM IPTG at 16 °C for 16 h resulted in robust expression relative to uninduced controls. WT and K301M CD2AP were purified to near homogeneity using single-step Ni–NTA affinity chromatography. This streamlined purification strategy differed from previously reported multistep protocols and yielded intact, full-length CD2AP suitable for downstream biophysical characterization (**Figures 1A and 1 B**)^23^ (M&M).

**Figure 1.**
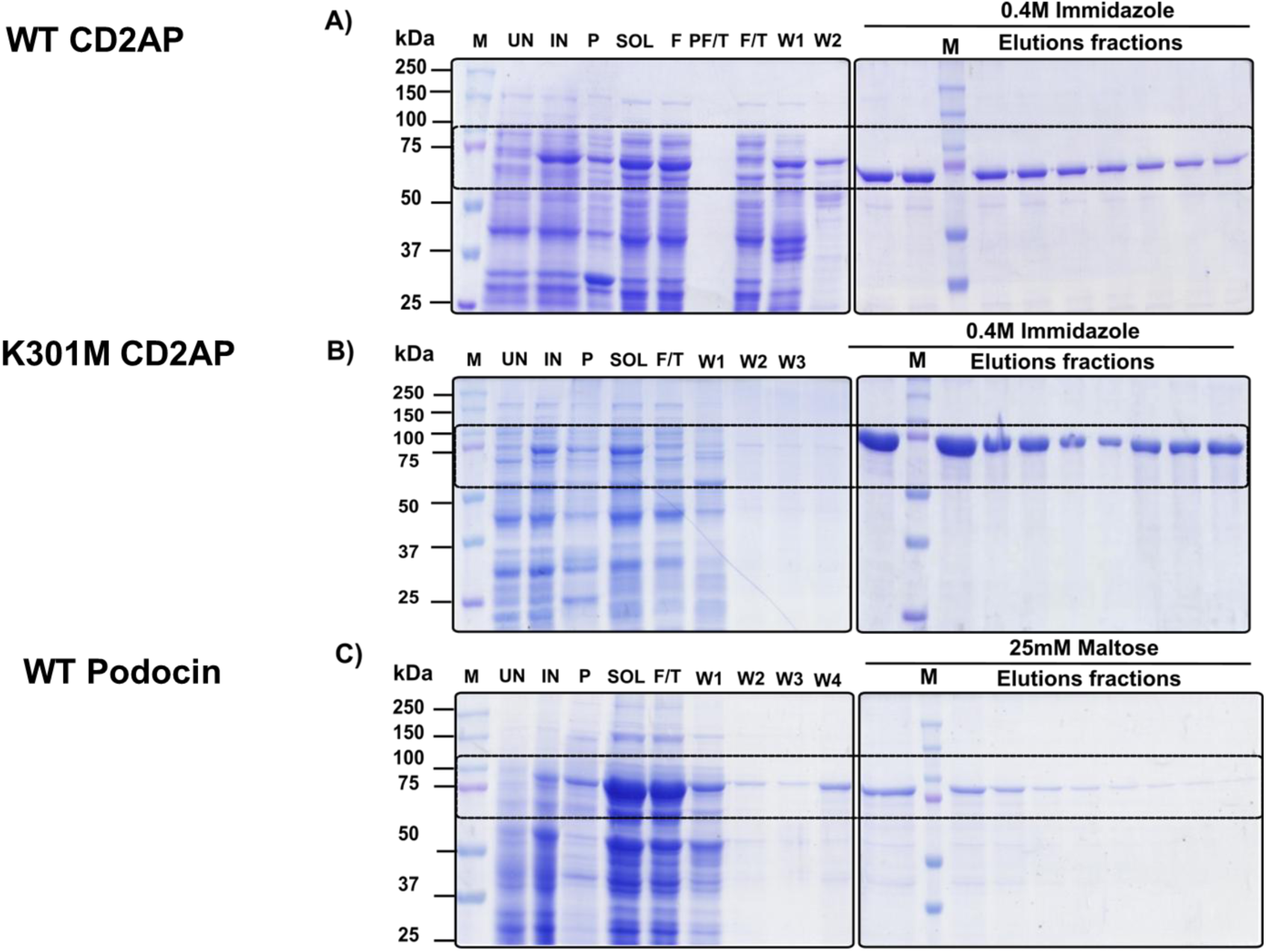
Recombinant production of WT and K301M CD2AP and Podocin: **(A)** SDS–PAGE analysis showing efficient expression and Ni–NTA purification of wild-type (WT) CD2AP, with enrichment of the full-length ∼73 kDa protein in soluble and imidazole-eluted fractions. **(B)** K301M CD2AP displays comparable expression, solubility, and purity, with no detectable degradation. **(C)** SDS–PAGE of MBP–Podocin purified by amylose affinity chromatography, revealing a dominant ∼82 kDa species and minimal contaminants, indicating suitability for downstream interaction and biophysical analyses.

Recombinant human Podocin was expressed in *E. coli* Arctic cells as an N-terminal maltose-binding protein (MBP) fusion. It was predominantly recovered in the soluble fraction following cell lysis and clarification. Purification by amylose affinity chromatography yielded a major species migrating at approximately 82 kDa on SDS–PAGE, consistent with the predicted molecular mass of the MBP–Podocin fusion protein (MBP, ∼42 kDa; Podocin, ∼40 kDa). The stepwise maltose elution profile indicated efficient retention of the fusion protein on the column, with the majority of Podocin recovered in later elution fractions. SDS–PAGE analysis confirmed high purity of the preparation with minimal contaminating host proteins (**Figure 1C**). The solubility and integrity of the purified protein indicate that the MBP tag facilitated proper folding and recovery of Podocin, yielding material of sufficient quality for subsequent structural and biophysical analyses.

### K301M substitution perturbs oligomeric architecture

K301M substitution perturbs the oligomeric architecture of CD2AP. Lys301 is located within the SH3-C region and proximal to the coiled-coil segment that stabilizes CD2AP homo-oligomerization. Substituting the positively charged lysine with a hydrophobic methionine would likely disrupt electrostatic contacts and local packing, thereby compromising higher-order assembly. To assess the biophysical consequences of this substitution, we generated the K301M variant by overlap PCR (Supplementary Figure 1). We purified both WT and mutant proteins using a previously optimized one-step affinity protocol with minior modifications (M&M) (**Figure 1A**, B) ^23^. SEC analysis revealed that WT CD2AP eluted as a heterogeneous ensemble, with broad overlapping peaks corresponding to approximately 12-mer (∼869 kDa) and 9-mer (∼669 kDa) assemblies. In Contrast, K301M produced well-resolved peaks at approximately 15.5 mL and 16.5 mL, consistent with lower-order oligomers, predominantly hexamers (∼440 kDa) and trimers (∼200 kDa) (**Figure 2 A&B**). SDS PAGE of SEC fractions confirmed protein integrity, excluding proteolysis as a contributor to later-eluting species (Supplementary Figure 2A, B). Overlaying chromatograms revealed a marked rightward shift for the K301M (**Figure 2C**), and comparison with molecular-weight standards verified its confinement to lower oligomeric ranges (**Figure 2D**).

**Figure 2.**
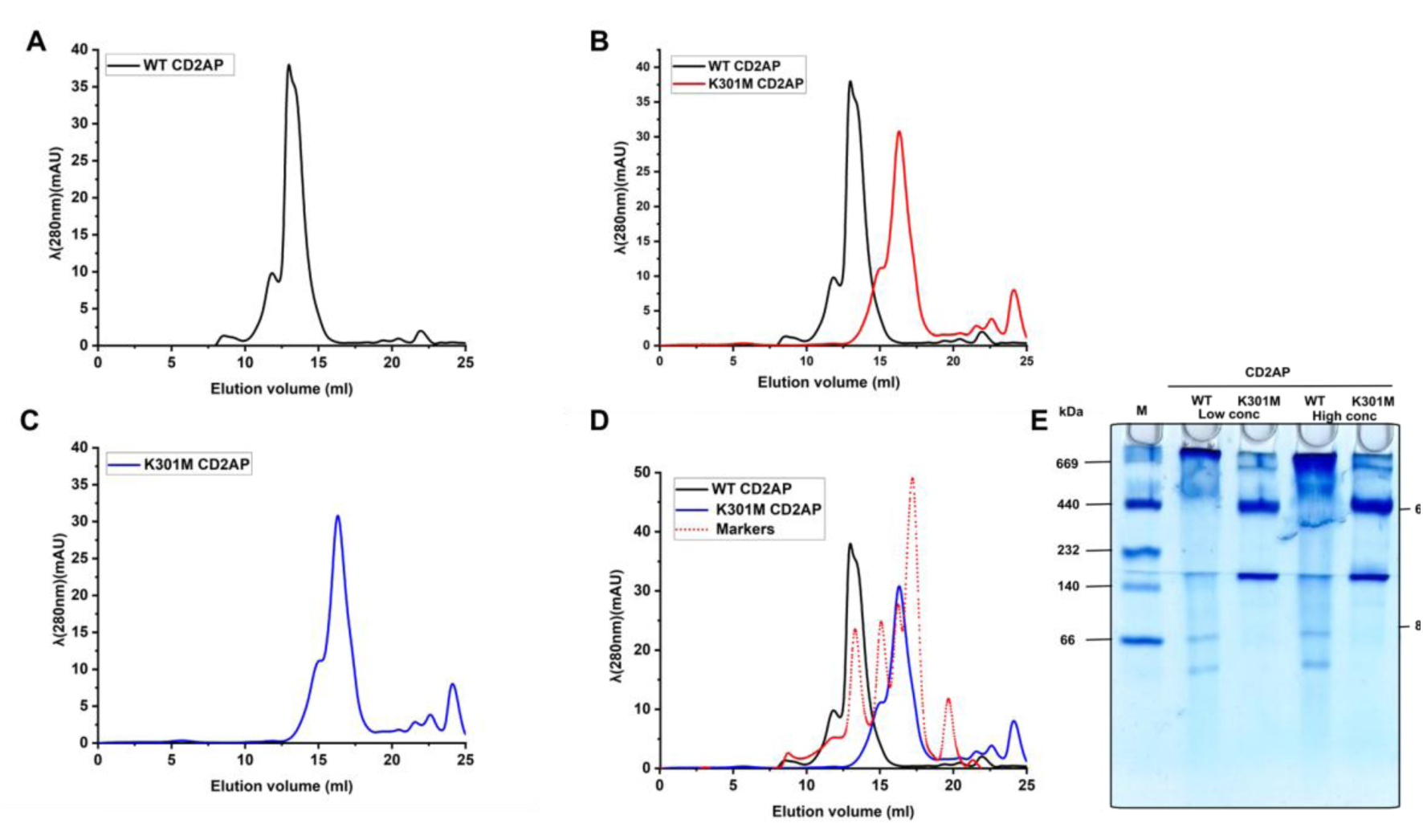
K301M substitution perturbs oligomeric architecture: **(A, B)** Size-exclusion chromatography profiles show that WT CD2AP elutes as higher-order oligomeric assemblies, whereas K301M shifts to later elution volumes consistent with smaller oligomers. **(C)** Overlay of WT and mutant traces highlight a pronounced rightward shift for K301M. **(D)** Comparison with molecular-weight standards confirms that K301M is confined to lower oligomeric ranges. **(E)** Blue native PAGE corroborates SEC results, revealing stable higher-order multimers for WT and restriction of K301M to lower-order species, indicating impaired self-association.

Further BN-PAGE analysis corroborated these observations, revealing apparent differences in the oligomeric organization of affinity-purified WT and K301M CD2AP.WT CD2AP resolved predominantly as 12-mer (∼869 kDa) and 9-mer (∼669 kDa) species, with a substantial fraction retained at the top of the gel, consistent with the presence of larger, higher-order assemblies. In Contrast, K301M displayed clear bands corresponding to hexamers (∼440 kDa) and trimers (∼200 kDa), aligning with hexameric and trimeric molecular-weight markers. When protein concentration was increased, WT CD2AP showed progressive enrichment of higher-order multimers, whereas K301M remained confined to its lower-order oligomeric states, with no evidence of larger complex formation, even under high-loading conditions (**Figure 2E**). Together, SEC and BN-PAGE analysis indicate that WT CD2AP forms a dynamic continuum of higher-order oligomers, whereas the K301M substitution restricts assembly to smaller, sharply defined complexes. These findings identify Lys301 as a critical determinant of coiled-coil packing stability and self-association, demonstrating that its substitution markedly compromises CD2AP’s ability to form and stabilize its native oligomeric architecture.

### Dynamic light scattering reveals conformational expansion of K301M CD2AP

Dynamic light scattering (DLS) further probed the hydrodynamic and conformational properties of CD2AP variants. Under native conditions, WT CD2AP exhibited a hydrodynamic radius (Rh) of ∼20.9 nm with a polydispersity index (PDI) of 26.3%, consistent with a heterogeneous population of higher-order oligomers. In Contrast, the K301M mutant displayed an increased Rh (∼34.1 nm) and higher PDI (∼29.1%) (**Figure 3(A)**. Upon denaturation with 8 M urea, both WT and K301M proteins underwent dramatic expansion (Rh ∼135.5 nm) with sharply reduced PDI values (<2%), indicative of complete unfolding and formation of a uniform monomeric ensemble (**Figure 3(B)**. Thus, while WT and K301M CD2AP differ markedly in their native oligomerization, both converge to similar unfolded states under denaturing conditions.

**Figure 3.**
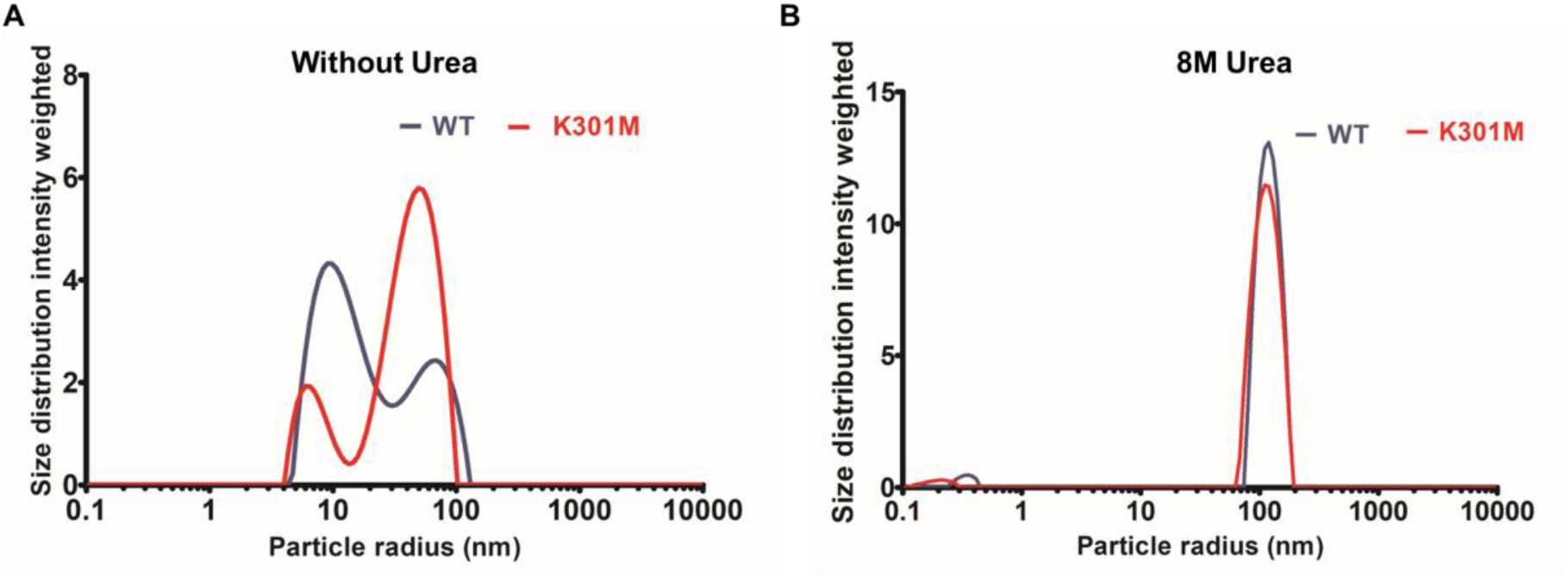
Dynamic light scattering reveals conformational expansion of K301M CD2AP: **(A)** Under native conditions, WT CD2AP exhibits a heterogeneous population consistent with higher-order assemblies, whereas K301M displays an increased hydrodynamic radius, indicative of conformational expansion despite reduced oligomeric order. **(B)** In 8 M urea, both proteins exhibit similar hydrodynamic profiles, indicating a complete loss of native quaternary structure and comparable denatured-state behavior.

### K301M Disrupts Secondary Structure and Tertiary Packing in CD2AP

The oligomerization differences observed in SEC and BN-PAGE provide a structural context for interpreting the CD spectra of WT and K301M CD2AP. These architectural distinctions align closely with the CD results. Despite both proteins exhibiting the characteristic far-UV signature of a largely disordered structure, except for the coiled coil and SH3 domains, K301M displayed reduced ellipticity at 208–215 nm, consistent with the loss of backbone regularity expected for a compact, conformationally restricted oligomer (**Figure 4 A, B, &C**). The divergence became even more pronounced in near-UV CD, where WT showed well-ordered aromatic packing, whereas K301M exhibited diminished and inverted ellipticity, reflecting the disruption of the long-range tertiary contacts that are stabilized in WT’s oligomeric assemblies. Thus, the CD signatures directly mirror the oligomerization patterns. WT’s higher-order assemblies support more ordered secondary and tertiary structural features. Furthermore, in the case of K301M, the mutation collapses this architecture into lower oligomers with an altered secondary and tertiary structural organization (**Figures 4A’, B’&C’**).

**Figure 4.**
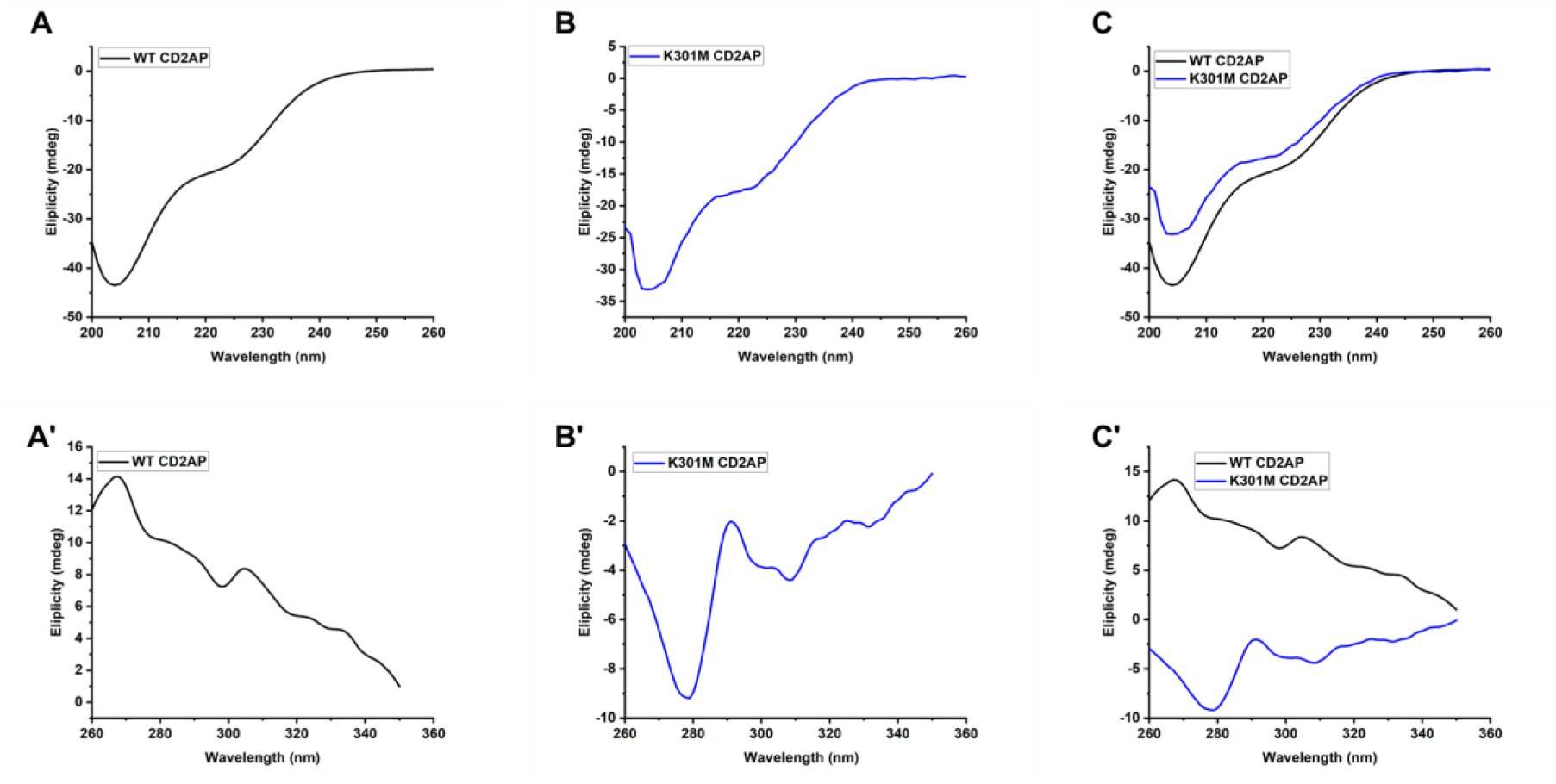
K301M disrupts secondary structure and tertiary packing: **(A–C)** Far-UV circular dichroism spectra reveal that both WT and K301M CD2AP retain a largely disordered, coil-rich fold, with reduced ellipticity in the mutant indicating altered secondary-structure organization. **(A′–C′)** Near-UV CD spectra show well-defined aromatic environments in WT, whereas K301M exhibits diminished and inverted signals, reflecting pronounced disruption of long-range tertiary packing.

### K301M Triggers Early Secondary-Structure Collapse Under Thermal Stress

To assess the thermal stability of CD2AP’s secondary-structure elements within its oligomeric assemblies, we performed far-UV CD thermal melts from 5–95 ^ο^C. WT CD2AP maintained its characteristic minima at 208 nm and 222 nm across a broad temperature range, showing only gradual reductions in ellipticity (**Figure 5A**). Further analyzing the effect of temperature on secondary structures at 208, 212, 218, and 222 nm, corresponding to Beta-sheet, and helical contributions, respectively, revealed minimal changes up to ∼65 °C, followed by a moderate decline at higher temperatures without complete loss of signal (**Figure 5B**). These data indicate that WT CD2AP retains substantial secondary-structure content despite thermal perturbation, consistent with a distributed, non-cooperative unfolding mechanism expected for an extended, partially disordered 12-/9-mer scaffold. In Contrast, the K301M variant exhibited markedly reduced thermal resilience. Far-UV CD spectra showed an earlier onset of structural decay, with pronounced loss of ellipticity, particularly at 208 nm, beginning at 40–50 °C (**Figure 5A′**). The wavelength-resolved melt profiles confirmed a steeper decrease in signal intensity across all monitored bands (**Figure 5B′**). Unlike WT, K301M failed to maintain stable minima at elevated temperatures, and its partial signal recovery at 80–95 °C was muted, indicating compromised secondary structure integrity and a diminished capacity to withstand thermal stress. Together, these data demonstrate that while both proteins preserve the overall coil-rich/disordered fold characteristic of CD2AP, the K301M mutation significantly weakens local secondary-structure packing and lowers the threshold for thermal disruption. This heightened sensitivity aligns with the mutant’s restricted oligomeric state (hexamers/trimers) and supports a model in which Lys301 is a key determinant of the structural stability of CD2AP’s multivalent assemblies.

**Figure 5.**
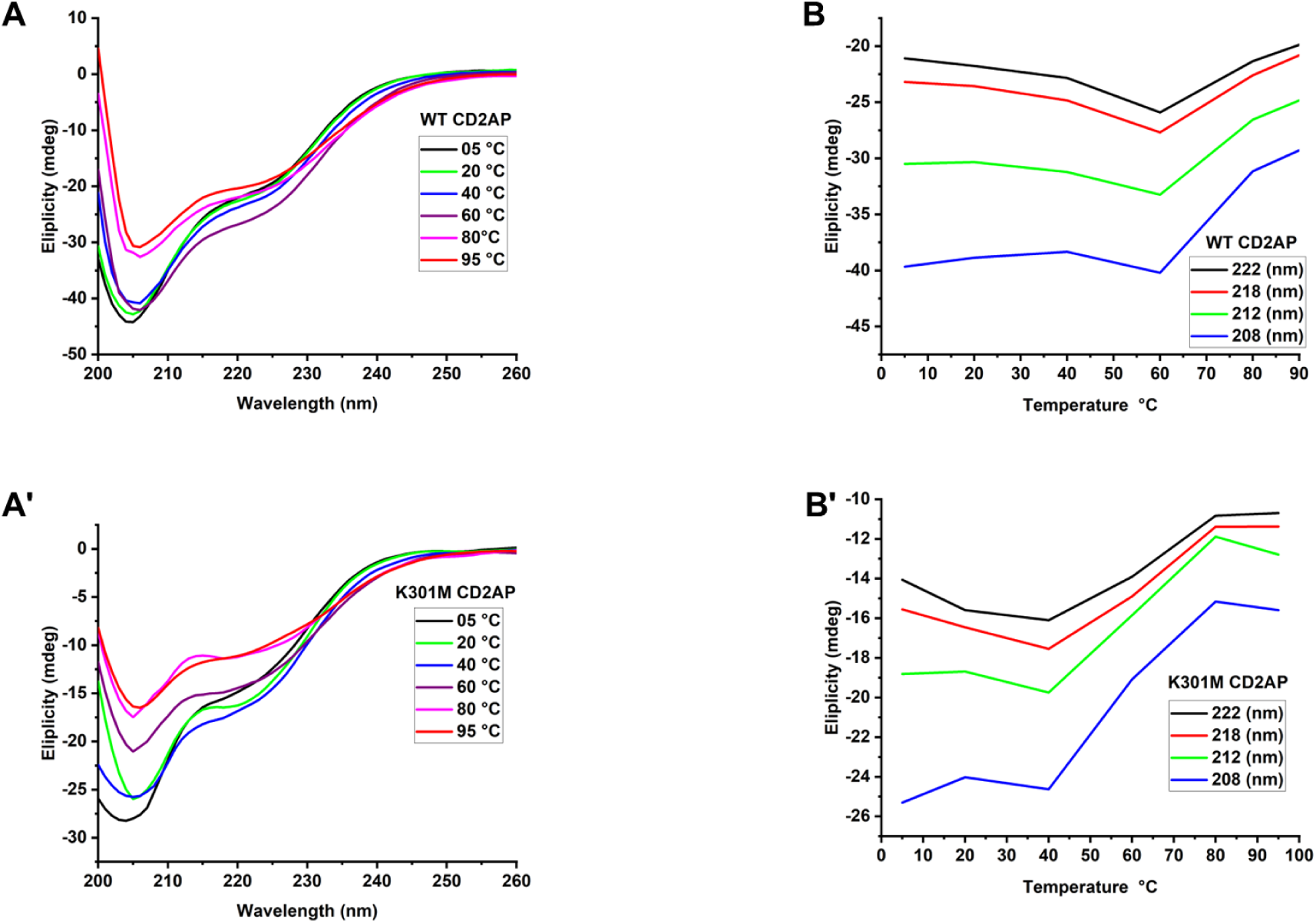
K301M lowers the thermal resilience of CD2AP secondary structure: **(A, B)** WT CD2AP maintains characteristic far-UV CD minima across a broad temperature range, consistent with distributed, non-cooperative unfolding. **(A′,B′)** K301M shows earlier and steeper loss of ellipticity, indicating reduced thermal stability and weakened secondary-structure packing relative to WT.

### K301M mutation accelerates tertiary destabilization of CD2AP

Temperature-dependent tryptophan fluorescence revealed distinct stability profiles for WT and K301M CD2AP. WT exhibited high fluorescence intensity at low temperatures (5–20°C), followed by a progressive decline between 20°C and 60°C (**Figure 6A**). The steep drop in intensity at ∼40–60°C corresponds to dissociation of the 12-mer/9-mer scaffold. In comparison, the shallow slope at higher temperatures (>60°C) indicates slow tertiary destabilization of dissociated subunits (**Figure 6B**). This stepwise, non-cooperative trajectory reflects the distributed unfolding typical of partially disordered multimers. K301M displayed an altered trajectory. Fluorescence began to decrease at much lower temperatures (20–40°C), exhibiting a rapid early decline consistent with the collapse of hexmeric and trimeric oligomers (**Figure 6A′**). The 341 nm λ max intensity slope was steeper than WT and reached near-baseline by ∼60°C (**Figure 6B′**). The lack of a transition plateau indicates that K301M undergoes direct oligomeric collapse coupled to tertiary destabilization, consistent with impaired quaternary packing and reduced structural buffering capacity relative to WT. Cooling spectra confirmed minimal refolding in both proteins; however, K301M retained greater structural distortion at all temperatures (**Figure 6C and C′**).

**Figure 6.**
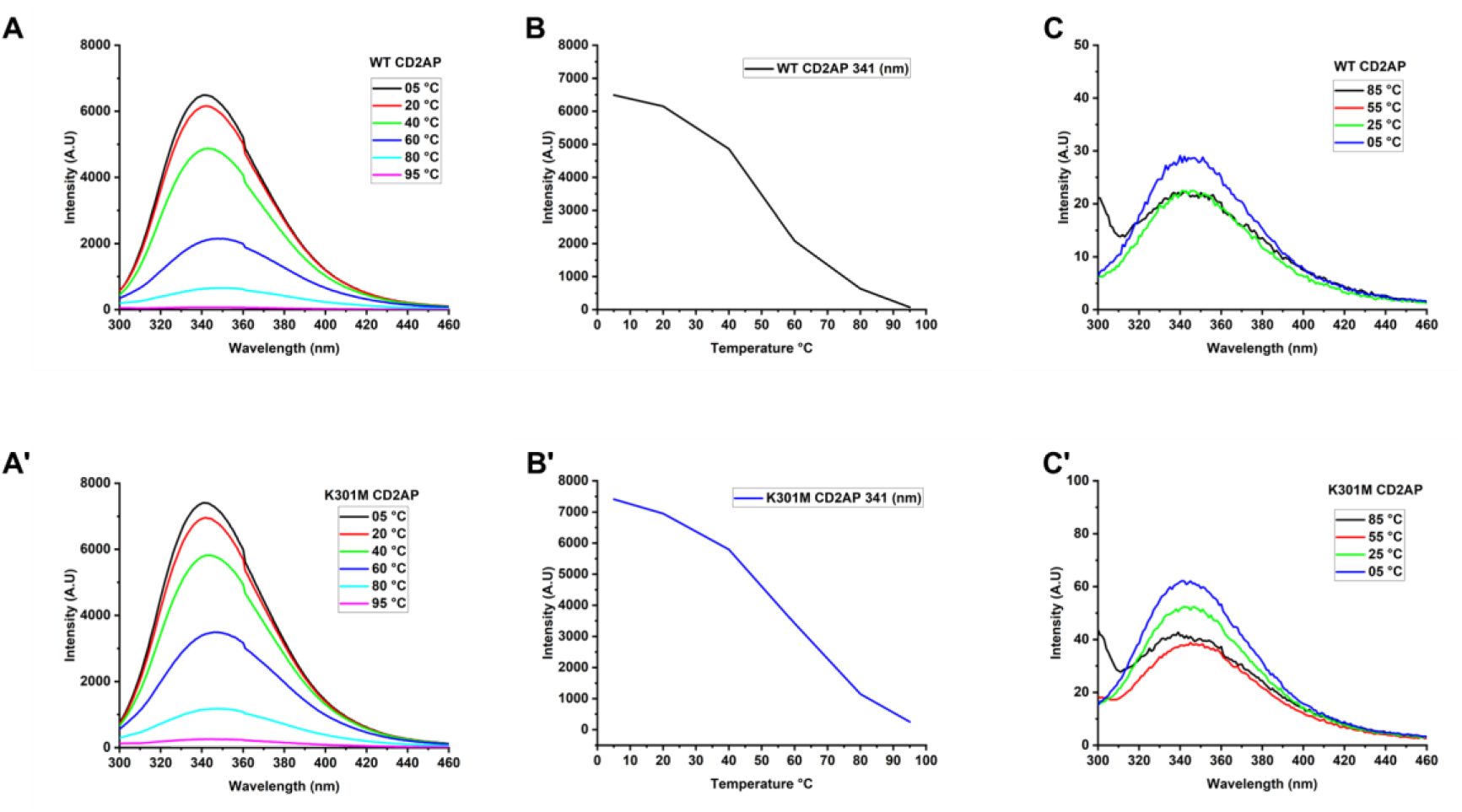
K301M accelerates tertiary destabilization and limits refolding: **(A, B)** The intrinsic tryptophan fluorescence of WT CD2AP decreases gradually with increasing temperature, consistent with progressive tertiary destabilization. **(C)** Cooling spectra reveal limited refolding. **(A′–C′)** K301M exhibits earlier fluorescence loss, a steeper unfolding trajectory, and minimal recovery upon cooling, indicating compromised tertiary stability and reduced structural buffering.

### SAXS defines distinct solution architectures of WT and K301M CD2AP

Small-angle X-ray scattering (SAXS) was used to determine the solution conformations of WT and K301M CD2AP. The WT scattering curve showed higher overall intensity and a characteristic downward slope on the log–log plot, indicative of extended, higher-order assemblies (**Figure 7A**). Guinier analysis yielded an increased radius of gyration (Rg 10.7 nm), consistent with a broad distribution of elongated multimeric species (**Figure 7B**). The corresponding pair-distance distribution function P(r) exhibited an asymmetric profile with a long tail and a maximum dimension (Dmax) of 35.5 nm, supporting the presence of extended oligomers, consistent with the known 9-mer and 12-mer assemblies (**Figure 7C**). By Contrast, the K301M mutant displayed a distinct scattering pattern with reduced intensity and a steeper decay, characteristic of more compact particles (**Figure 7A**’). The Guinier region yielded a smaller Rg (9.6 nm), and the P(r) distribution was narrower with a reduced Dmax of 33.9 nm, indicative of lower-order assemblies such as trimers and hexamers (**Figure 7B’&C’**). The well-defined linear Guinier region in the mutant further supported loss of extended multimers and a shift toward compact, less elongated conformations. Ab initio SAXS envelopes calculated for both proteins were superimposed with AlphaFold-generated multimeric models, showing excellent agreement between the experimental envelopes and the predicted WT higher-order assemblies, whereas the K301M envelope aligned only with compact, lower-order models (**Figure 7D&D’**). Together, SAXS analysis demonstrates that WT CD2AP self-associates into extended, higher-order oligomeric assemblies in solution, whereas the K301M substitution disrupts this architecture, favoring compact and lower-order species. These structural changes are consistent with the reduced stability and impaired cooperative behavior observed in CD spectroscopy and intrinsic fluorescence analyses.

**Figure 7.**
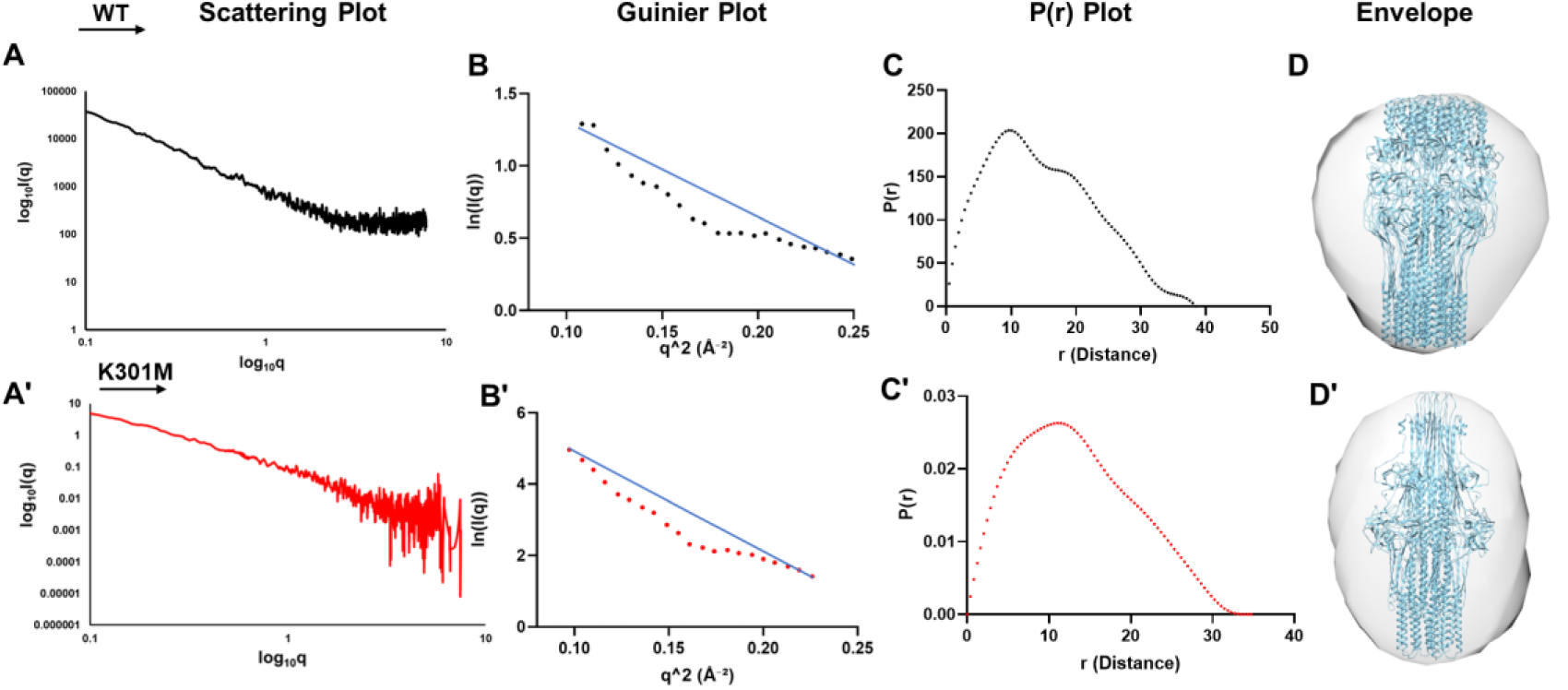
SAXS reveals distinct solution architectures of WT and K301M CD2AP: **(A, A′)** The scattering profiles indicate extended, higher-order assemblies for WT and more compact particles for K301M. **(B, B′)** Guinier analysis reveals a reduced radius of gyration in the mutant. **(C, C′)** Pair-distance distribution functions show a broader, elongated profile for WT and a narrower distribution for K301M. **(D, D′)** Ab initio SAXS envelopes align with higher-order AlphaFold models for WT and compact oligomers for K301M, confirming mutation-induced architectural collapse.

### K301M mutation disrupts the CD2AP–podocin complex

Previous studies have established that CD2AP interacts with Podocin, and the K301M mutation may compromise this interaction ^4, 28, 30–32^. In vitro pulldown assays using purified recombinant proteins further assessed the CD2AP–Podocin interaction. WT CD2AP exhibited robust binding to Podocin, whereas the K301M variant demonstrated markedly reduced binding. Ni–NTA resin controls confirmed the interaction specificity, ruling out nonspecific resin adherence (**Figure 8A and B**). These findings support the hypothesis that the K301 residue contributes to the stabilization of CD2AP–Podocin complexes, suggesting that its mutation perturbs the proper assembly of SD Protein complexes.

**Figure 8.**
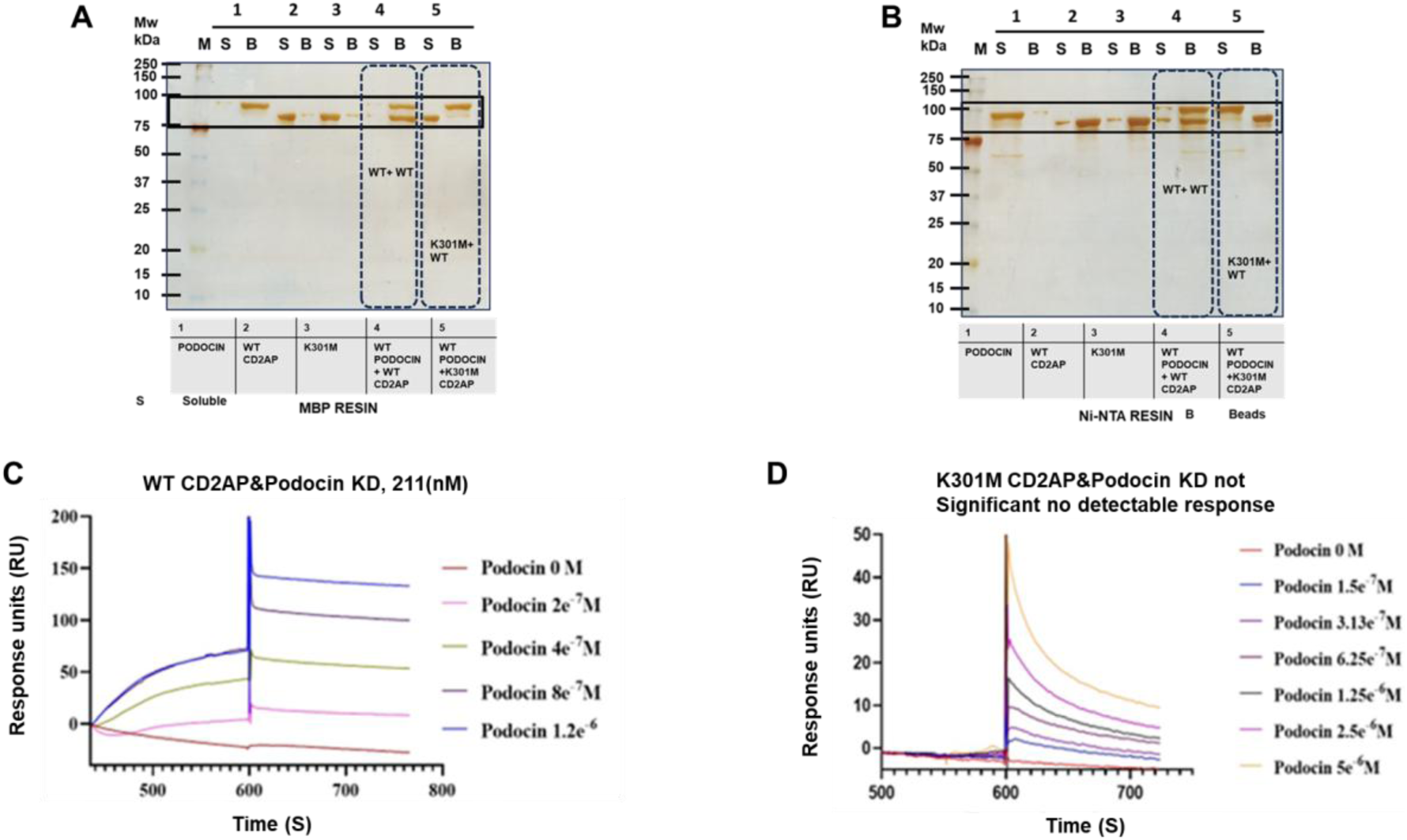
K301M mutation abolishes CD2AP–podocin interaction: **(A, B)** In vitro pulldown assays demonstrate robust association between WT CD2AP and Podocin, whereas K301M fails to retain Podocin under reciprocal binding configurations. **(C)** Surface plasmon resonance shows concentration-dependent binding of Podocin to WT CD2AP with measurable affinity. **(D)** K301M exhibits no detectable binding, confirming loss of podocin recognition.

### K301M mutation abolishes CD2AP–podocin interaction

Podocin produced clear, concentration-dependent association and dissociation phases when flowed over CD2AP-coated surfaces, yielding a dissociation constant of ∼211 nM, consistent with strong affinity binding. By Contrast, K301M CD2AP generated only minimal response units across all tested concentrations, displaying no measurable association and no definable KD. The absence of detectable binding indicates a complete loss of affinity for Podocin (**Figures 8C&D**). Together, the pulldown and SPR analyses show that Lys301 is essential for stabilizing the CD2AP–podocin interface. The K301M mutation eliminates detectable binding, reinforcing the notion that disruption of higher-order CD2AP assembly, as demonstrated by SEC and BN-PAGE, directly impairs its ability to engage Podocin.

### Mutation-dependent remodelling of the CD2AP interactome

Proteomic profiling reveals mutation-dependent remodeling of the CD2AP interactome. To assess the impact of the disease-associated K301M substitution on CD2AP-mediated protein interactions, affinity pulldown assays coupled with LC–MS/MS analysis were performed using recombinant His-tagged WT and K301M CD2AP proteins (**Figure 9A**). Purified proteins were immobilized on Ni–NTA beads and incubated with human podocyte cell-free lysates to enrich for interacting partners. Bound protein complexes were resolved by SDS–PAGE (**Figure 9B**) and subjected to mass spectrometric identification. Comparative proteomic analysis demonstrated distinct yet partially overlapping interaction profiles among WT CD2AP, K301M CD2AP, and bead-only control samples (**Figure 9C**). Across all conditions, 533 proteins were detected, reflecting shared and nonspecific binders. The control sample contained 345 proteins, of which 137 overlapped with the mutant pulldown, indicating increased nonspecific association in the K301M interactome.

**Figure 9.**
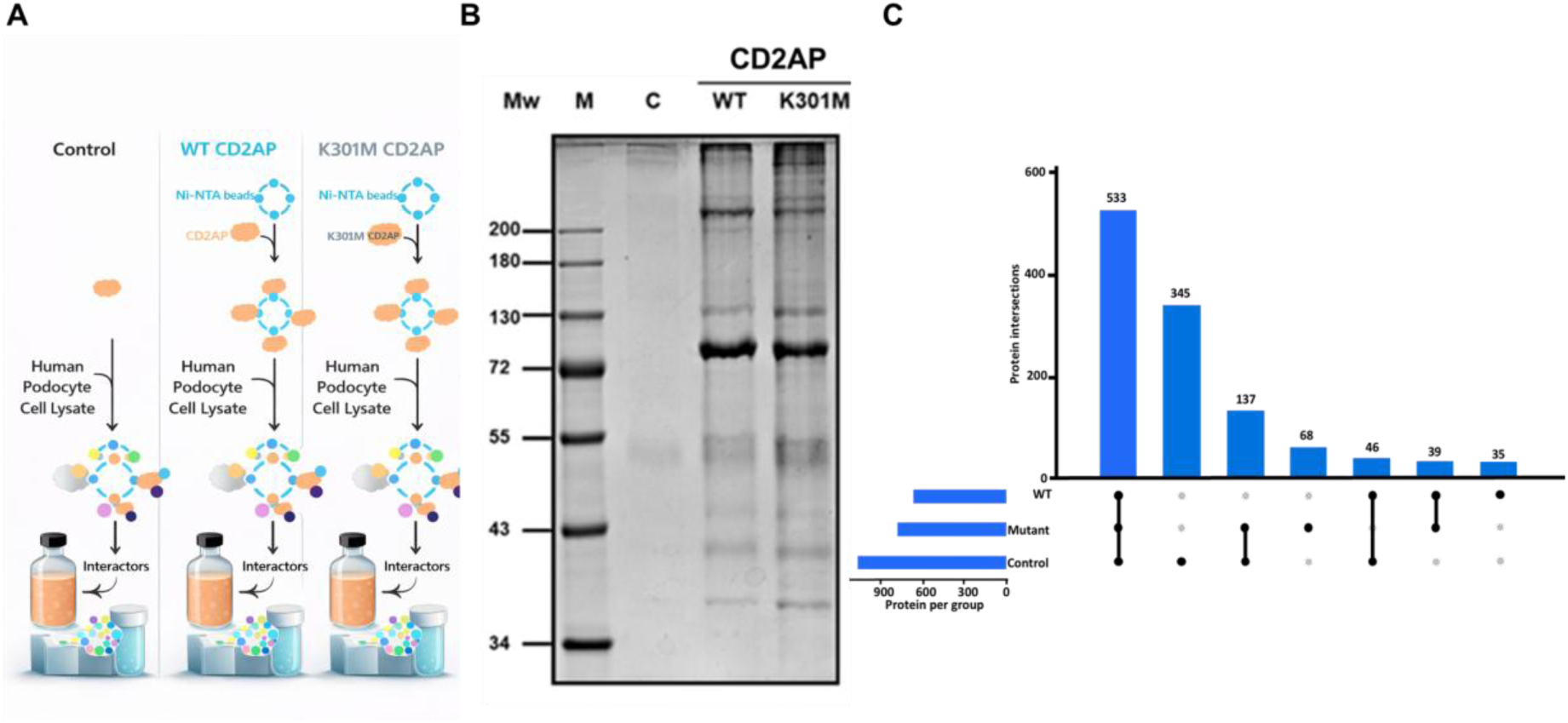
Mutation-dependent remodelling of the CD2AP interactome: **(A)** Schematic of affinity enrichment and proteomic workflow. (B) SDS–PAGE of affinity-purified complexes from podocyte lysates. **(C)** Upset plot comparing interactomes reveals that WT CD2AP engages a broader interaction network, whereas K301M shows reduced and redistributed binding partners, indicating impaired interaction specificity.

In Contrast, WT CD2AP recovered 37 proteins that were not detected in either the mutant or control samples, whereas the K301M pulldown identified 68 proteins unique to the mutant condition. A subset of proteins was shared between WT and mutant CD2AP, suggesting partial preservation of interaction capacity despite the mutation. Notably, the K301M substitution was associated with a marked redistribution of CD2AP binding partners. As detailed in Supplementary Table 1, WT-specific interactors were enriched for RNA-binding proteins, ribosomal components, and scaffold-associated factors. In Contrast, the K301M variant exhibited increased association with metabolic and mitochondrial proteins. The loss of WT-specific interactors accompanied by the gain of mutant-selective associations reflects mutation-induced disruption of CD2AP structure and oligomeric organization. This interactome reprogramming is predicted to impair CD2AP-dependent scaffolding functions, which are essential for slit diaphragm integrity and podocyte signaling. Together, these findings identify altered interaction specificity as a molecular basis for slit diaphragm disorganization and podocyte dysfunction in CD2AP-associated glomerular disease.

## Discussion

The SD is a specialized adherens-type junction between podocyte foot processes that forms the final barrier preventing urinary protein loss ^6–9^. Structurally, it represents a supramolecular network composed of key interacting proteins, including nephrin, Podocin, CD2AP, ZO-1, TRPC6, NEPH1–3, and others ^10, 11, 33–38^. The functional integrity of this macromolecular lattice depends on precise organization and stable association among its constituents ^10, 11, 18, 33–37, 39–48^. Preliminary studies on CD2AP focused mainly on its role in signal transduction. Studies by Huber et al. showed that CD2AP, upon interacting with nephrin and Podocin, facilitated nephrin signaling and initiated PI3K/AKT-dependent signal transduction in podocytes ^12^. It is revealed that nephrin interacts with the C-terminal domain of CD2AP, and CD2AP, in turn, interacts with Podocin and the actin cytoskeleton of podocytes, thus acting as an adaptor protein ^40, 44^. It is, therefore, understandable that mutations in CD2AP lead to severe proteinuria, as seen in mice with missense and frameshift mutations in CD2AP that developed heavy proteinuria and died after a few weeks ^49^ ^50^. Although these studies demonstrate a central role for CD2AP at the SD, studies detailing the structural features of CD2AP were limited to the crystal structures of SH3 domains ^51^. It was reported that SH3 domains have a natural affinity for proline-rich motifs, which coincidentally are present in the C-terminal region of Podocin ^52^. Despite its pivotal role, the structural determinants governing CD2AP function and the molecular mechanisms by which monogenic mutations (K301M) disrupt the integrity and fishnet architecture of SD remain incompletely understood.

Comprehensive biophysical, structural, and proteomic analyses demonstrate that the K301M substitution induces conformational perturbations and structural plasticity in CD2AP, impairing its oligomerization and remodeling its interaction network. Our results showed that (WT) CD2AP self-assembles into large oligomeric complexes (∼9–12 subunits), whereas the K301M variant forms smaller, less stable assemblies (∼3–6-mers). SAXS confirmed that WT CD2AP adopts an extended, flexible conformation consistent with its role as a multivalent scaffold capable of bridging nephrin, Podocin, and other SD components into higher-order complexes. In Contrast, the K301M mutant exhibits a more compact and rigid conformation, indicating a loss of structural plasticity and reduced multivalency, which are crucial for cooperative network formation at the SD ^53^. In the WT CD2AP, the assembly into higher-order oligomers (∼9–12-mers) expands its interactome, enabling multivalent contacts with key SD proteins such as nephrin and Podocin, as well as cytoskeletal adaptors. The extensive CD2AP interaction network forms a stable, fishnet-like lattice that maintains narrow and regulated filtration pores, preventing albumin loss. The K301M mutant, confined to smaller oligomeric assemblies (∼3–6-mers), displays a contracted interactome and reduced multivalent binding capacity. The consequent loss of cooperative interfaces likely disrupts crosslinking among SD components and weakens their anchorage to the actin cytoskeleton, compromising lattice tension and pore selectivity. This conformational and interaction remodeling impairs the recruitment and stabilization of SD complexes, leading to loss of filtration fidelity and proteinuria characteristic of glomerular disease.

Consistent with earlier predictions that 65–75% of CD2AP is intrinsically disordered ^18^, our far-UV CD spectra confirm that both WT and K301M retain a largely coil-rich architecture dominated by a strong minimum near 205 nm. Such extensive intrinsic disorder is characteristic of signaling adaptors, where flexible regions mediate multivalent electrostatic interactions and embed short molecular recognition features (MoRFs) capable of disorder-to-order transitions upon partner engagement ^54–58^. These dynamic segments likely enable CD2AP to orchestrate SD signaling and to stabilize interaction networks with key partners, such as podocin ^54, 57, 59–62^. Although the K301M mutation does not abolish this global disordered state, our biophysical analyses reveal that it markedly compromises the structural resilience and conformational adaptability of CD2AP.

These structural perturbations propagate directly to CD2AP oligomeric assembly. SEC and BN-PAGE analyses demonstrate that wild-type (WT) CD2AP assembles into higher-order multimers (9–12-mers), whereas the K301M variant is confined to lower-order trimers and hexamers, indicating a failure to generate the multivalent scaffolds required for SD integrity and signaling. This architectural collapse has clear functional consequences: WT CD2AP engages Podocin with moderate affinity (KD ≈ 221 nM), while K301M shows no detectable binding, manifested as non-saturable SPR sensorgrams and markedly reduced retention in pulldown assays. These findings identify Lys301 as a critical structural and electrostatic determinant that couple’s oligomer stability to productive podocin recognition.

To define how these molecular defects translate into podocyte dysfunction, we compared the interaction landscapes of WT and K301M CD2AP by affinity purification–mass spectrometry. WT CD2AP formed a coherent interactome enriched in scaffold-associated and RNA-binding proteins essential for SD organization and signaling. In contrast, the K301M mutation triggered extensive interactome remodeling, characterized by loss of core partners and aberrant association with noncanonical metabolic proteins, indicative of impaired interaction specificity. Integrating these proteomic insights with our biophysical and structural data, we propose that Lys301 is central to the formation of conformationally flexible, multivalent CD2AP assemblies that stabilize SD archit33ecture. Disruption of this assembly logic abolishes podocin engagement, rewires interaction networks, and provides a mechanistic framework for how CD2AP variants precipitate slit diaphragm failure and proteinuric kidney disease ^54, 57, 59–61, 63, 64,62^.

in conclusion, our integrative structural and biophysical analysis delineates the structural logic underlying CD2AP function and elucidates how the K301M mutation mechanistically disrupts its role at the SD. By coupling intrinsic disorder with controlled oligomerization, CD2AP achieves both flexibility and strength attributes critical for the dynamic yet stable filtration barrier. We are continuing our quest to reveal high-resolution studies integrating cryo-EM and time-resolved biophysics to understand how SD is assembled as a molecular sieve to filter the proteins and minor molecular perturbations results in proteinuric kidney disease.

## Data Sharing Statement

SAXS datasets generated in this study have been deposited in the Small Angle Scattering Data Bank (SASDB) under accession codes SASDXJ6 (wild-type CD2AP) and SASDXK6 (K301M CD2AP). Mass spectrometry data supporting the findings of this study are available from the corresponding authors upon reasonable request. All other data are included within the article and its supplementary information files.

## Acknowledgments

**Author Contributions: AHQ and AKP** conceived and designed the study. **AHQ** performed cloning, protein expression, purification, biophysical, and structural experiments, and analyzed the data. **RMT and RR** contributed to SAXS data collection, analysis, and interpretation. **IF** assisted with protein–protein interaction assays and mass spectrometry–based proteomic experiments. **SM** contributed to spectroscopic measurements and data analysis. **ERR** provided expertise in data interpretation and critical methodological input. **AHQ**, **AKP**, and **IF** drafted the manuscript. All authors reviewed, edited, and approved the final version of the manuscript.

## Funding Information and Acknowledgments

Acknowledgments: PAK acknowledges IoE grants of University of Hyderabad (RC1-20-21). PAK acknowledges DST-FIST and DBT-BUILDER Infrastructure at the Department of Biochemistry and School of Life Sciences, University of Hyderabad. Authors thank Hira Singh Gariya (CDRI) for assistance with SPR.

## Statement

During the preparation of this work, the authors used ChatGPT (OpenAI) to support editing of grammar, sentence structure, and overall readability. Further verified by the authors, who take full responsibility for the final manuscript.

---

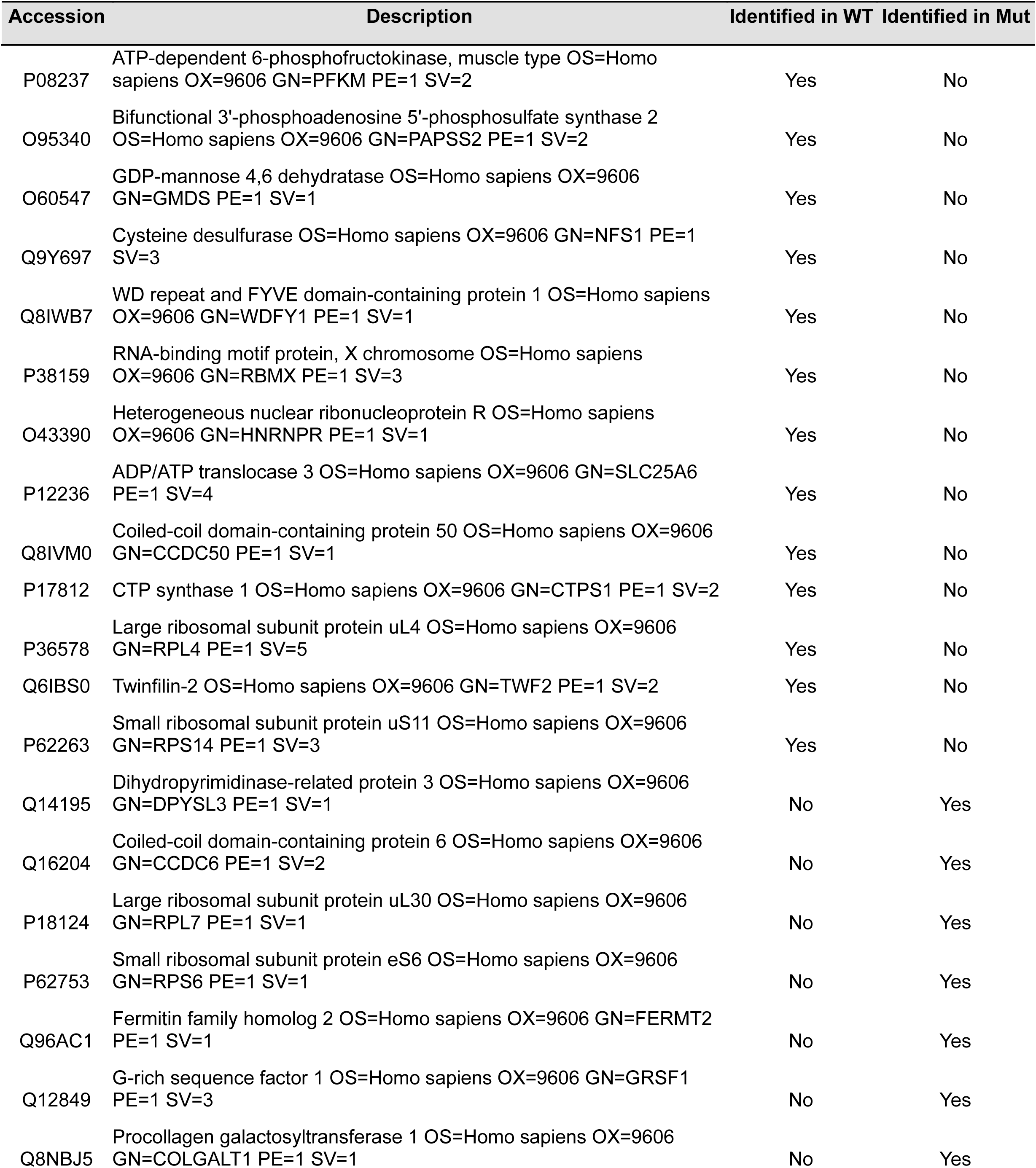
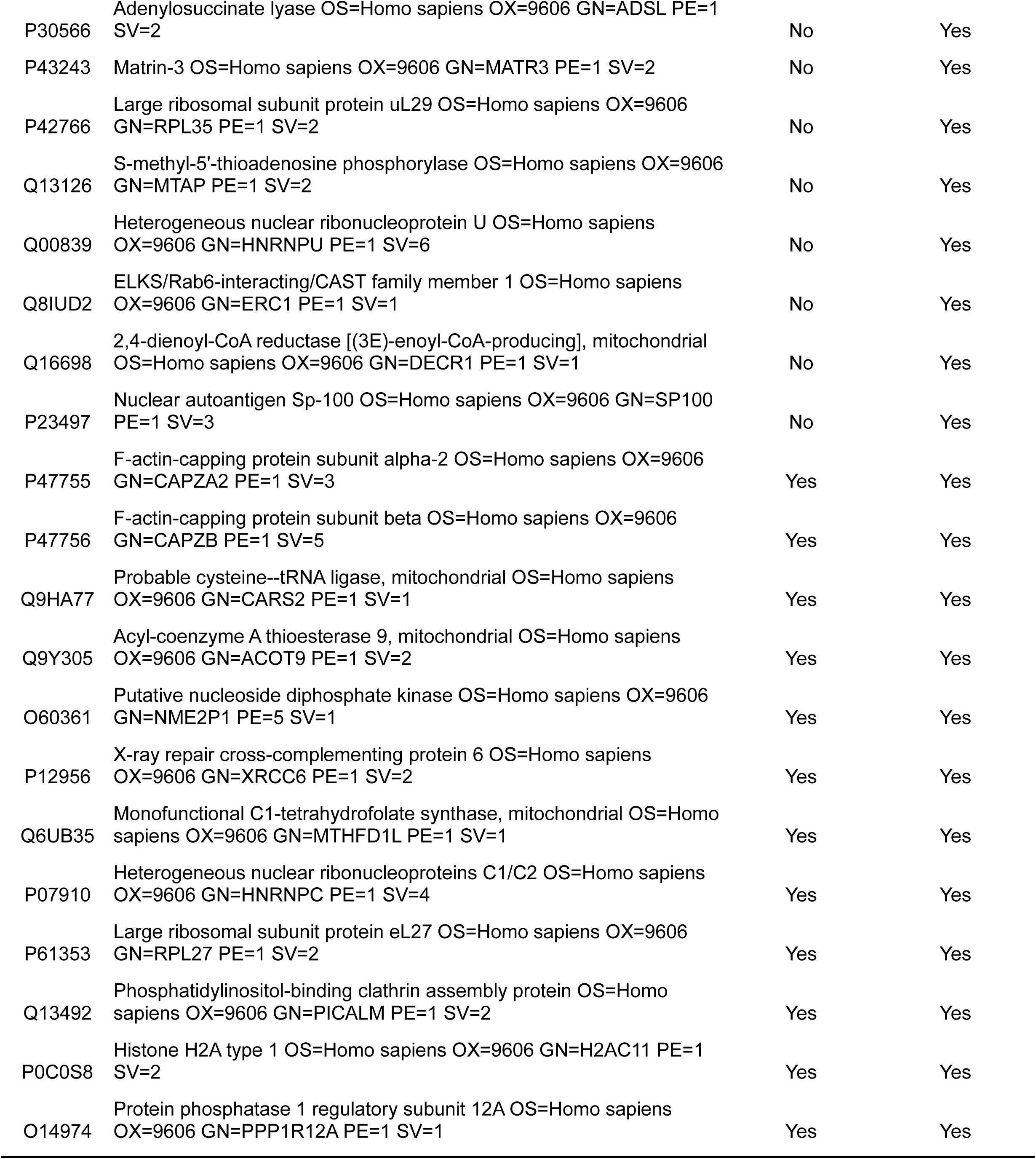

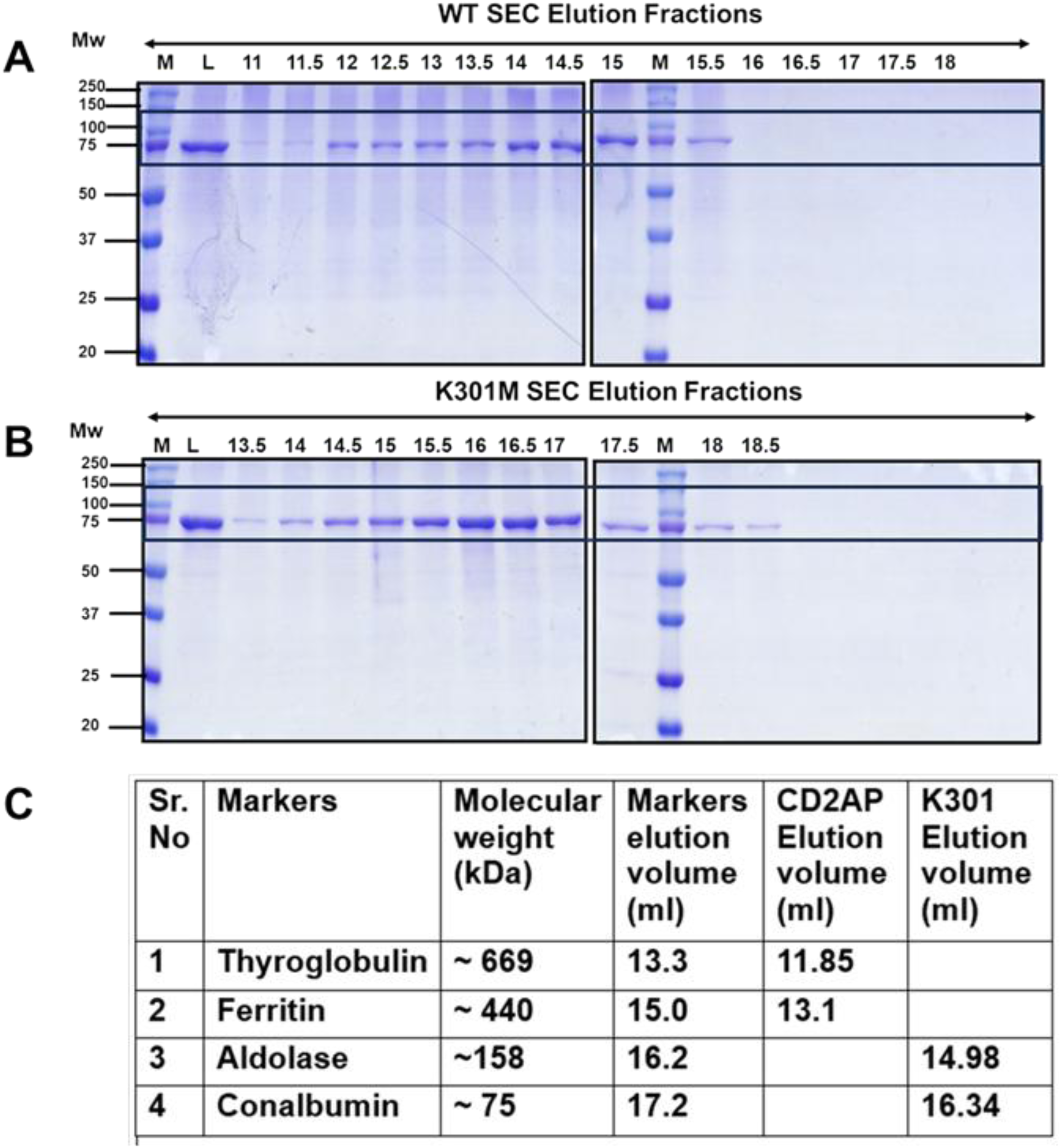

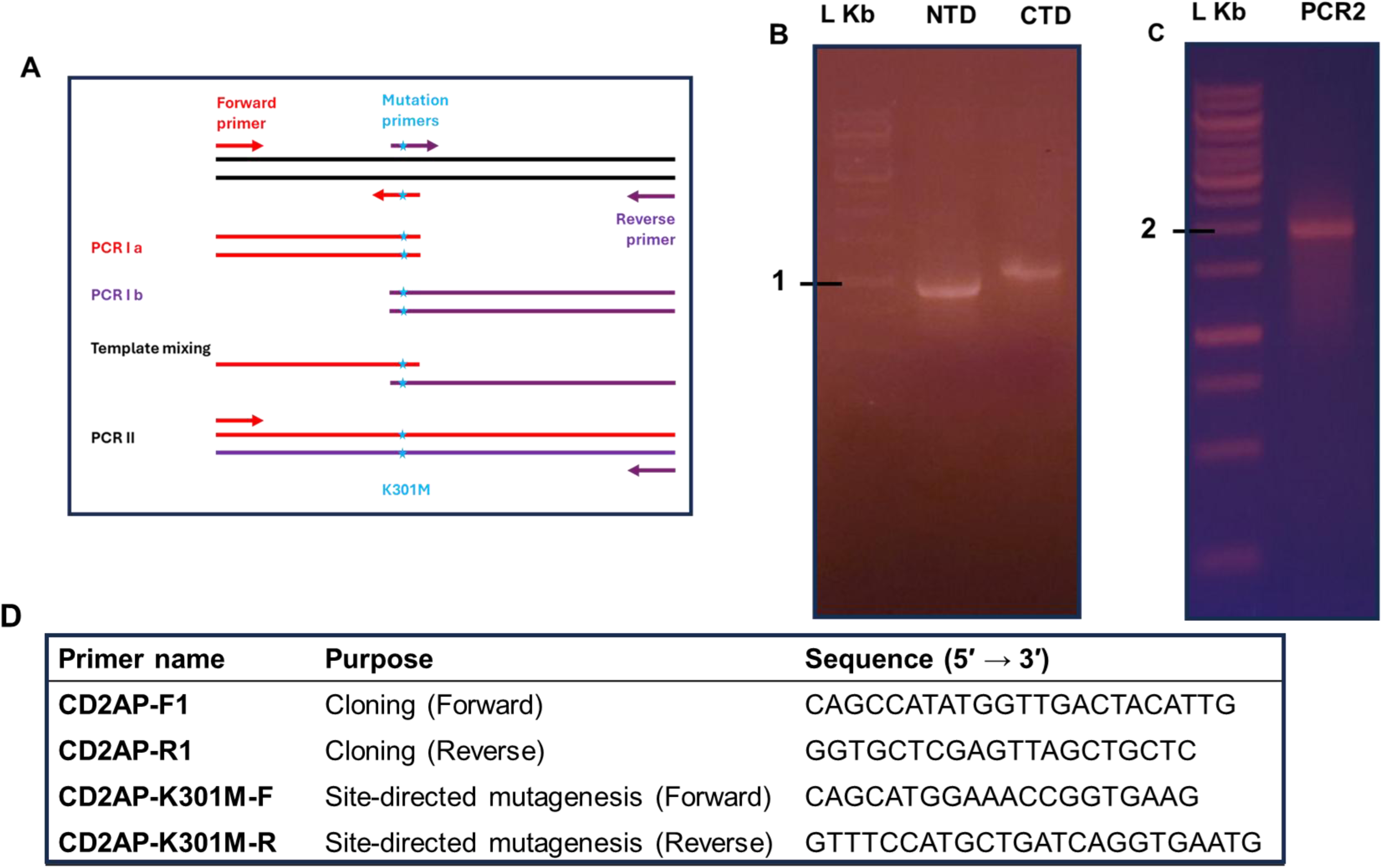

